# Diminishing neuronal acidification by channelrhodopsins with low proton conduction

**DOI:** 10.1101/2023.02.07.527404

**Authors:** Rebecca Frank Hayward, F. Phil Brooks, Shang Yang, Shiqiang Gao, Adam E Cohen

## Abstract

Many channelrhodopsins are permeable to protons. We found that in neurons, activation of a high-current channelrhodopsin, CheRiff, led to significant acidification, with faster acidification in the dendrites than in the soma. Experiments with patterned optogenetic stimulation in monolayers of HEK cells established that the acidification was due to proton transport through the opsin, rather than through other voltage-dependent channels. We identified and characterized two opsins which showed large photocurrents, but small proton permeability, PsCatCh2.0 and ChR2-3M. PsCatCh2.0 showed excellent response kinetics and was also spectrally compatible with simultaneous voltage imaging with QuasAr6a. Stimulation-evoked acidification is a possible source of disruptions to cell health in scientific and prospective therapeutic applications of optogenetics. Channelrhodopsins with low proton permeability are a promising strategy for avoiding these problems.

**Statement of Significance:** Acidification is an undesirable artifact of optogenetic stimulation. Low proton-permeability opsins minimize this artifact while still allowing robust optogenetic control.

## Introduction

Channelrhodopsins are light-gated ion channels that are widely used to modulate the activity of neurons and other excitable cells.^1^ In addition to research use, these tools are entering clinical practice as a treatment for forms blindness^2,3^ and are under consideration for treatments of other disorders of neural excitability.^4–6^ Every ion channel carries current through one or more ions, and so the induced change in membrane voltage is always accompanied by a change in ionic concentrations. In both research and clinical applications, one must consider whether the ionic changes in the cell have effects beyond the purely electrical effects of the channel.

Ionic perturbations are largest when (a) the basal intracellular concentration of the relevant ion is low, (b) the surface-to-volume ratio is high and (c) the channel is activated chronically. Sodium, potassium, and chloride ion concentrations in cells are typically in the millimolar range, and thus the fractional changes in concentration of these ions due to channel opening are typically < 1%. In contrast, protons and calcium ions have low free concentrations, typically ^∼^100 nM or lower. In these cases, the ionic fluxes due to channel opening can substantially perturb the concentration.

The amount of charge flow required to change membrane voltage by a given amount is proportional to the capacitance, and hence the membrane surface area. This charge is diluted into the volume. For this reason, ionic concentrations in small structures with high surface-to-volume ratios are more labile to optogenetic perturbations than are concentrations in large structures. One must therefore consider whether optogenetic tools substantially perturb ionic concentrations in thin structures such as axons, dendrites, or dendritic spines. Finally, ionic concentrations equilibrate much more slowly than does membrane potential. Brief optogenetic stimuli may have negligible effects on concentrations, whereas chronic stimuli with high duty cycle may have cumulative effects. Concerns about long-term consequences of optogenetic stimulation are particularly relevant to prospective therapeutic applications, where the tools may be used over long times in patients.

Channelrhodopsins have been shown to acidify cells, while light-driven outward proton pumps, such as Archaerhodopsin 3, alkalize cells.^7,8^ Indeed, optogenetically triggered alkalization has been proposed as a tool to control cell death.^9^ Intracellular acidification can suppress neuronal excitability and also suppress vesicle release,^10–12^ but has also been reported to enhance release of adenosine^13,14^ and dopamine.^15^ Changes in intracellular pH can also affect cell differentiation^16^ and metabolism^17^ and survival.^7,8,11,18^ Nominally similar channelrhodopsins have been reported to evoke opposite behavioral effects in live mice^19^. The cause of these differences is not known, but it is possible that these differences could be due, in part, to differences in ionic selectivity.

For these reasons, it is important to quantify and ultimately minimize perturbations to cellular pH from optogenetic tools. We combined channelrhodopsin stimulation with a red-shifted fluorescent pH sensor, pHoran4,^20^ for measurement of pH changes during optogenetic stimulation. We used patterned optogenetic stimulation in gap junction-coupled cellular monolayers to establish that the acidification was due to proton flux directly through the channelrhodopsin. We then tested two new opsins, ChR2-3M and PsCatCh2.0,^21^ with a low proton permeability and found minimal perturbations to cellular pH. We performed a detailed electrophysiological and photophysical characterization of these opsins and showed that they are compatible with simultaneous voltage imaging. The new opsins may be promising for clinical applications where acidification is undesirable.

## Results

### CheRiff acidifies neurons

We developed lentiviral constructs and optical stimulus protocols for simultaneous optogenetic stimulation and pH measurements (Fig. 1A). For the actuator we used CheRiff-GFP, a non-selective cation channel with an activation peak at 460 nm.^25^ For the pH measurement we used pHoran4,^20^ a red-shifted reporter with an excitation peak at 547 nm and a pK_a_ of 7.5. We calibrated the pH response of pHoran4 in permeabilized HEK293T (HEK) cells and in cultured neurons (Methods; Figure 1 – figure supplement 1) and then used this calibration to convert changes in fluorescence to changes in pH. We assumed an initial pH of 7.3 for all the cells (Fig. 1B). Since our measurements focused on relative pH changes as opposed to absolute pH, modest deviations from this assumption would not affect the interpretation of the following results.

**Figure 1.**
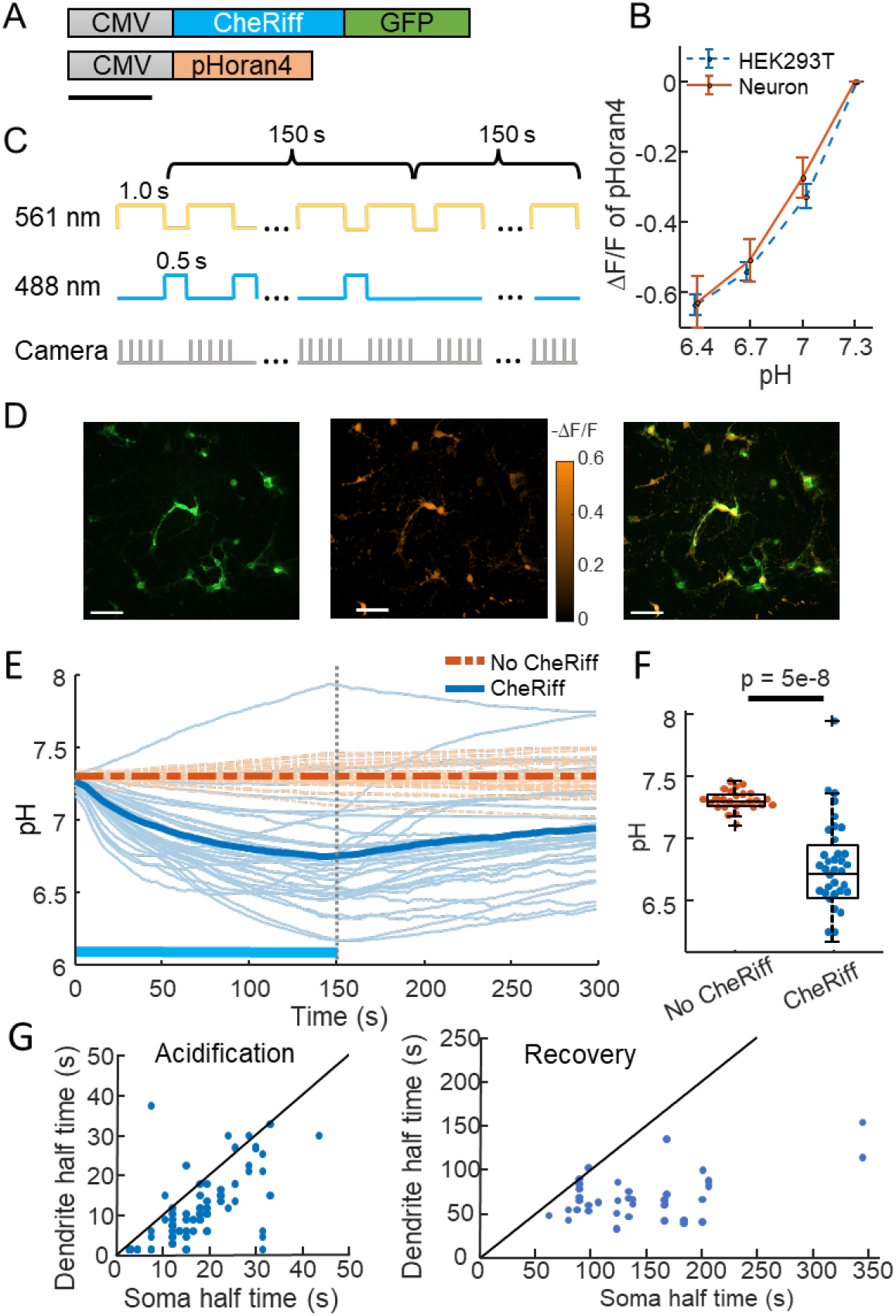
CheRiff acidifies polarized cells. A) Genetic constructs for simultaneous optogenetic stimulation and pH imaging. B) Calibration of pHoran4 pH sensor in HEK cells and neurons. Error bars represent S.D. of *n =* 8 measurements in HEK cells, 42 measurements in neurons. C) Protocol for measuring pH responses to optogenetic stimulation. Stimulation (blue) and measurement (yellow) were interleaved for 150 s; then pH recovery was measured for 150 s without optogenetic stimulation. D) Example images of cultured neurons showing (left) GFP fluorescence, a marker for CheRiff expression, (middle) -ΔF/F in the pHoran4 channel after 150 s of the protocol shown in (C), (right) merge. Scale bars 100 µm. E) Time-course of pH in cultured neurons. Cells expressing pHoran4 but not CheRiff did not acidify. Bold lines show population average. F) CheRiff-expressing neurons acidified to a pH of 6.76 ± 0.35 (mean ± S.D., *n* = 34 cells). Neurons not expressing CheRiff had significantly less acidification, pH = 7.3 ± 0.08 (mean ± S.D., *n* = 26 cells, *p* = 5e-8 Wilcoxon rank sum test). Box plots show inter-quartile ranges, tick-marks show data range, + shows outlier. G) Half-time of (left) acidification or (right) recovery for neuron somas vs dendrites stimulated with the protocol in (C). Black line shows equal kinetics.

We imaged pH changes in cultured neurons during CheRiff stimulation. We alternated epochs of optogenetic stimulation (488 nm, 400-800 mW cm^-2^, 0.5 s) and pH imaging (561 nm, 100-200 mW cm^-2^, 1 s) to avoid crosstalk of the blue light into the pH recordings (Fig. 1C). The pH dynamics were much slower than 1.5 s, so this process did not sacrifice information.

After 150 s of stimulation and imaging, the pH had decreased within the neurons that expressed CheRiff-GFP, from 7.3 to 6.76 ± 0.35 (mean ± S.D., *n* = 34 cells, Fig. 1D, E), corresponding to an approximately 3-fold increase in concentration of free protons. We then measured the pH recovery for an additional 150 s without optogenetic stimulation (Fig. 1E). Recovery was slow, returning to only pH 6.95 ± 0.33 (mean ± S.D.) after 150 s. Control experiments in neurons that expressed pHoran4 but not CheRiff showed a small fluorescence increase during the stimulation period (0.11 ΔF/F), much smaller in magnitude and opposite in sign compared to the change in cells expressing CheRiff and pHoran4 (-0.34 ΔF/F). We attribute the slight increase in fluorescence of the CheRiff-negative cells to a blue light-mediated photo-activation artifact, as has been seen in other fluorescent reporters.^26^ The mean photo-artifact from CheRiff-negative cells was subtracted from all recordings of CheRiff-positive cells prior to analysis.

We observed that stimulation-induced pH changes and post-stimulation recovery were faster in the dendrites than in the soma (Fig. 1G). This effect is most likely due to the higher surface-to-volume ratio of thin processes. For a given proton current density across the membrane, the change in local proton concentration is greater in a thin tube than in the large soma.

### Acidification is via proton transport through the opsin

We next sought to determine to what extent the acidification was due to proton transport through the opsin vs through depolarization-induced opening of endogenous proton-permeable channels or other activity-dependent acidification mechanisms (e.g. metabolic shifts). Working in HEK cells, we expressed CheRiff, pHoran4 and a doxycycline-inducible inward rectifying potassium channel, K_ir_2.1, (Methods, Fig. 2A). The K_ir_2.1 channel polarized the HEK cells to a resting potential of approximately -70 mV,^27^ providing a driving force for proton entry. K_ir_2.1 has been reported not to carry a proton current itself.^28^ When the cells were grown into a confluent monolayer, they coupled electrically via endogenous gap junctions (Fig. 2B).^29^

**Figure 2.**
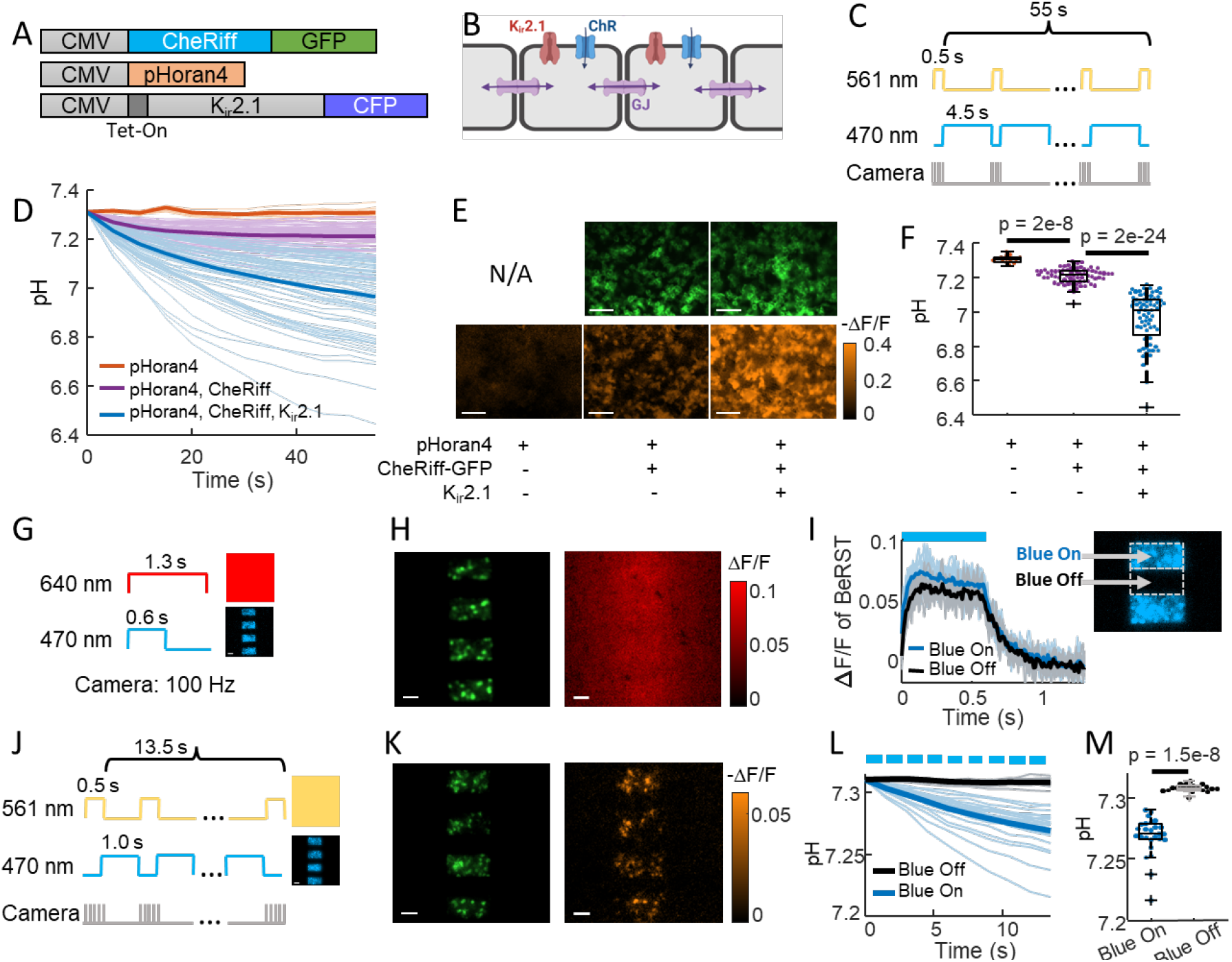
CheRiff exhibits high proton conductance. A) Genetic constructs for simultaneous optogenetic stimulation and pH imaging in polarized HEK293T cells. B) Diagram of HEK cell monolayer connected by gap junctions. C) Experimental paradigm for measuring pH responses to optogenetic stimulation. Stimulation (488 nm) and measurement (561 nm) were interleaved to avoid optical crosstalk. D) Time-course of pH in HEK cells. Expression of K_ir_2.1 increased the driving force for proton influx, substantially enhancing the acidification. E) Images of HEK cell monolayers showing (top) GFP fluorescence, a marker for CheRiff expression and (bottom) -ΔF/F in the pHoran4 channel after protocol sho©in (C). Scale bars 100 µm. F) Quantification of the data in (D-E). pHoran4 alone: pH = 7.31 ± 0.02 (mean ± S.D., *n* = 13 cells); CheRiff and pHoran4 pH = 7.21 ± 0.05 (*n* = 70 cells); CheRiff, pHoran4 and K_ir_2.1: pH = 6.96 ± 0.15 (*n* = 75 cells). Statistical comparisons via Wilcoxon signed rank test. G) Protocol for mapping voltage responses to patterned optogenetic stimulation (488 nm) via fluorescence of BeRST1 (640 nm exc.). H) Images of HEK cell monolayers showing (left) fluorescence of GFP with patterned blue illumination, (right) ΔF/F of BeRST1. Scale bars 100 µm. I) Time-course of BeRST1 fluorescence in HEK cells inside (Blue On) and outside (Blue Off) the optogenetic stimulus regions. J) Protocol for measuring pH responses to patterned optogenetic stimulation. Stimulation (488 nm) and measurement (561 nm) were interleaved to avoid optical crosstalk. K) (Left) Fluorescence of GFP with patterned blue illumination, (right) ΔF/F in the pHoran4 channel after protocol shown in (J). Scale bars 100 µm. L) Time-course of pH inside (Blue On) and outside (Blue Off) the optogenetic stimulus regions. M) Quantification of the data in (L). Directly stimulated cells acidified to pH = 7.27 ± 0.016 (mean ± S.D., *n* = 26 cells), indirectly depolarized cells (Blue Off) did not acidify: pH = 7.31 ± 0.003 (*n* = 19 cells, *p* = 1.5e-8 Wilcoxon signed-rank test).

We characterized these cultures using a wide-area “Firefly” microscope which provided spatially and temporally patterned illumination at 470 and 561 nm via a digital micromirror device (DMD).^30^ Fig. 2C shows the protocol for interleaved CheRiff stimulation and pH imaging. After 60 s of stimulation, cells expressing all three components were acidified to a pH of 6.96 ± 0.15 (mean ± S.D., *n* = 75 cells). Cells not expressing K_ir_2.1 had substantially less acidification (final pH 7.21 ± 0.05, mean ± S.D., *n* = 70 cells, *p* = 2e-24, Wilcoxon signed-rank test), confirming the importance of membrane voltage as a driving force for proton entry. Cells lacking both the CheRiff and the K_ir_2.1 did not show detectable acidification (final pH 7.31 ± 0.02, mean ± S.D., *n* = 13 cells, Fig. 2D-F), consistent with our results in neurons (Fig. 1E, F).

These results established that acidification occurred in non-excitable cells but left open the possibility that the HEK cells might contain an endogenous proton conductance that opened upon membrane depolarization. To test this possibility, we took advantage of the gap junctional coupling between cells in a confluent monolayer. Due to the gap junctions, local CheRiff activation led to depolarization of neighboring regions, with an electrotonic length constant of ^∼^300 µm.^29^ While protons can also diffuse through gap junctions, this process is orders of magnitude slower than propagation of membrane voltage.^31^ This difference in lateral electrical vs proton transport permitted us to indirectly depolarize cells via gap junction coupling, and to ask whether the acidification arose in all depolarized cells or only in cells with direct CheRiff activation.

We used the DMD to pattern the optogenetic stimulation into stripes, with a separation between the stripes of 95 µm, much smaller than the electrotonic length constant (Fig. 2G). We used the far-red voltage sensitive dye BeRST1^32^ to map the voltage changes throughout a monolayer of HEK cells expressing pHoran4, CheRiff, and K_ir_2.1. Fig. 2H shows the stimulus pattern (reported via fluorescence of CheRiff-GFP), and the electrical depolarization pattern. As expected from the strong gap junctional coupling, the optogenetically induced depolarization in the interstitial “Blue Off” regions was almost as large as in the directly stimulated “Blue On” regions (Fig. 2I).

We then mapped the pH changes over the whole field of view, using the same striped stimulation pattern alternating with wide-field yellow illumination for pH imaging (Fig. 2J). We expected that proton currents through CheRiff would follow the illumination pattern precisely, whereas proton currents through voltage-gated channels would follow the much smoother voltage profile. We quantified acidification after only a short (13.5 s) period of CheRiff stimulation to avoid possible confounding effects of lateral proton diffusion between cells. In the directly stimulated “Blue On” regions we observed robust acidification after 13.5 s (pH = 7.27 ± 0.016, mean ± S.D., *n* = 26 cells), and in the indirectly depolarized “Blue Off” regions we observed no acidification (pH = 7.31 ± 0.003, mean ± S.D., *n* = 19 cells, *p* = 1.5e-8 Wilcoxon signed-rank test, Fig. 2K-M). These results establish that CheRiff directly acidifies polarized cells via proton transport through the opsin, and that electrical depolarization alone is insufficient to drive acidification.

### ChR2-3M and PsCatCh2.0 are potent non-acidifying channelrhodopsins

Several channelrhodopsins were recently reported to have low proton conductivity,^21,33–36^ so we tested two of these for acidification in HEK cells and characterized their photophysical properties. PsCatCh2.0^21^ is derived from the highly blue shifted *Platymonas subcordiformis* PsChR^37^ via the L115C mutation and addition of a trafficking signal, ER export signal, and cleavable N-terminal Lucy-Rho signal peptide. Due to its high speed and high light sensitivity, this opsin has been used for visual function restoration in blind mice.^21^

The second opsin we tested is derived from the recently engineered ChR2-XXM (i.e. ChR2-D156H), which shows high photocurrent and high selectivity for Na^+^ and K^+^ over H^+^.^33,36^ Mutating H134 to Q at the intracellular gate further enhanced the Na^+^ and K^+^ selectivity (Fig. 3 – figure supplement 1) and photocurrent amplitude. Mutating E101 to N near the extracellular gate site also boosted the Na^+^ and K^+^ selectivity (Fig. 3 – figure supplement 1A) without affecting the photocurrent amplitude (Fig. 3 – figure supplement 1B). To optimize expression and trafficking, we added the same trafficking, ER export, and signal peptides as in PsCatCh2.0. We designate this triple mutant of ChR2 as “ChR2-3M”.

Following the same procedure as in Fig. 2C, we tested the acidification due to opsin stimulation in electrically polarized HEK cells expressing pHoran4, K_ir_2.1, and either CheRiff, ChR2-3M, or PsCatCh2.0 (Fig. 3A). After a 55 s stimulation and imaging protocol, CheRiff cells showed a decrease in pH as above (pH = 6.98 ± 0.15, mean ± S.D., n = 170 cells), while the new opsins did not show any significant changes in pH (ChR2-3M: pH = 7.31 ± 0.10, n = 63 cells; PsCatCh2.0: pH = 7.30 ± 0.03, n = 74 cells; p = 4e-31, p = 4e-35, Wilcoxon signed-rank test; Fig. 3B-D). These experiments confirmed the low proton permeability of ChR2-3M and PsCatCh2.0. In paired experiments, we used voltage imaging with BeRST1 to confirm that all three opsins induced depolarization in the HEK cell monolayers (Fig. 3 – figure supplement 2).

**Figure 3.**
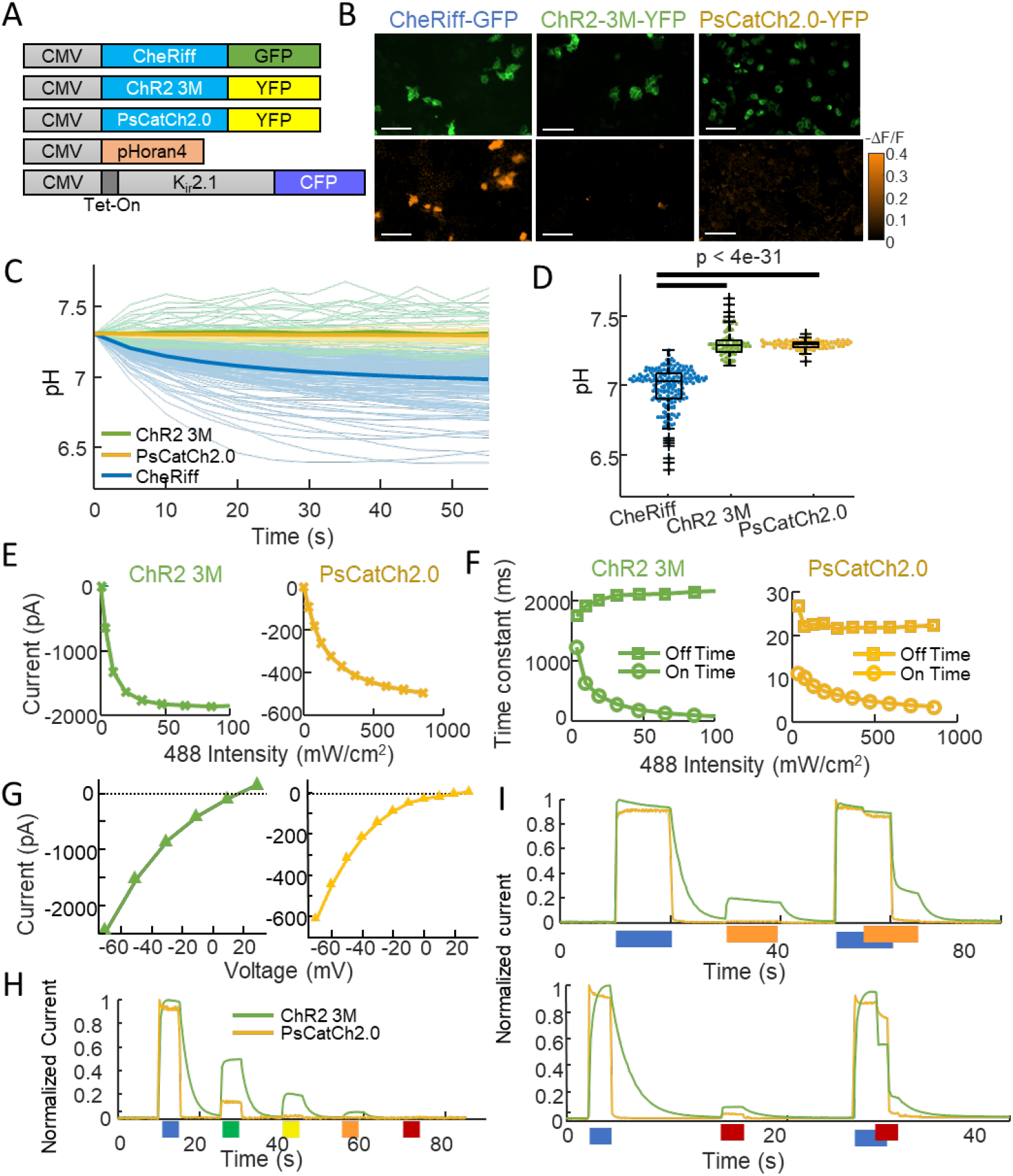
ChR2-3M and PsCatCh2.0 are potent non-acidifying channelrhodopsins. A) Genetic constructs for simultaneous optogenetic stimulation using channelrhodopsin variants and pH imaging in polarized HEK cells. B) Images of HEK cells showing (top) GFP or YFP fluorescence, a marker for channelrhodopsin expression and (bottom) -ΔF/F in the pHoran4 channel, measured after protocol shown in (2C). Scale bars 100 µm. C) Time-course of pH in HEK cells expressing the three opsins. D) Quantification of the©ta in (C). CheRiff: pH = 6.98 ± 0.15 (mean ± S.D., n = 170 cells); ChR2-3M: pH = 7.31 ± 0.10 (n = 63 cells); PsCatCh2.0: pH = 7.30 ± 0.03 (n = 74 cells); p = 4e-31, p = 4e-35, Wilcoxon signed-rank test. E-I) Whole-cell voltage clamp measurements on HEK cells expressing channelrhodopsins. E) Steady-state photocurrents as a function of blue illumination intensity. F) Opening and closing kinetics as a function of blue light intensity. G) Steady-state photocurrents as a function of holding voltage. H) Normalized photocurrents from stimulation with light at 488 nm, 532 nm, 561 nm, 594 nm, and 640 nm (50 mW cm^-2^ in all cases). I) Normalized photocurrents from combinations of blue (488 nm, 240 mW cm^-2^) and orange (594 nm, 1 W cm^-2^) or red (640 nm, 8 W cm^-2^) light corresponding to intensities typical for all-optical electrophysiology.

We then performed a detailed characterization of the new opsins using patch clamp electrophysiology in HEK cells (Fig. 3E-I, Table 1). Both opsins were sensitive to blue (488 nm) light. ChR2-3M passed unusually large steady-state photocurrents (1378 ± 618 pA, mean ± S.D., n = 4 cells) and was highly sensitive to blue light (Effective Power Density for 50% activation, EPD50 = 11.6 ± 8.7 mW cm^-2^, Fig. 3E), but had slow opening (τ_on_ = 57 ± 21 ms at saturating blue light) and very slow closing (τ_off_ = 1950 ± 500 ms, Fig. 3F). PsCatCh2.0 had somewhat smaller photocurrents (847 ± 359 pA, *n* = 6 cells), higher EPD50 (116 ± 13 mW cm^-2^), but very fast opening (τ_on_ = 4.2 ± 3.5 ms at saturating blue light) and closing (τ_off_ = 17.6 ± 3.4 ms).

**Table 1.** Comparison of channelrhodopsin gating properties. EPD50 is the effective power density for 50% activation. CheRiff data are from Ref. ^25^, Fig. S9 and Table S4. CheRiff reversal potential is from Ref. ^27^.

|  | Reversal Potential (mV) | t<sub>on</sub> fastest (ms) | t<sub>on</sub> at EPD50 (ms) | t<sub>off</sub> (ms) | EPD50 (mW cm<sup>-2</sup>) | Steady state photocurrent (pA) | I<sub>peak</sub>/I<sub>ss</sub> |
| --- | --- | --- | --- | --- | --- | --- | --- |
| **CheRiff** | 4 | 4.5 ± 0.3 | - | 16 ± 0.8 | 22 ± 4 | 1,300 ± 80 | 0.65 |
| **ChR2-3M (n=4)** | 16.6 ± 3.6 | 57 ± 21 | 800 ± 550 | 1,950 ± 500 | 11.6 ± 8.7 | 1,378 ± 618 | 1.00 |
| **PsCatCh2.0 (n=6)** | 12.3 ± 4.7 | 4.2 ± 3.5 | 9.3 ± 1.2 | 17.6 ± 3.4 | 116 ± 13 | 847 ± 359 | 0.92 |

Both opsins had positive reversal potentials (ChR2-3M: 16.6 ± 3.6 mV, PsCatCh2.0: 12.3 ± 4.7 mV), consistent with preferential Na^+^ and Ca^2+^ selectivity (Fig. 3G). A key attraction of PsCatCh2.0 for quantitative optogenetics experiments was its very flat-top response to a step in blue light. In contrast to CheRiff which shows substantial sag in photocurrent upon continuous illumination,^25^ PsCatCh2.0 showed almost no sag. This low extent of light induced inactivation appears to be, at least in part, a characteristic of this particular type of opsin from *P. subcordiformis*^37^.

For applications in all-optical electrophysiology (i.e. simultaneous stimulation and voltage or calcium imaging), it is critical that the light used for imaging a red-shifted reporter does not interfere with the action of the opsin. At 50 mW cm^-2^ excitation intensity, ChR2-3M retained substantial activation at 561 nm (20%) and 594 nm (4.8%), but undetectable activation at 640 nm (< 0.3%, Fig. 3H). PsCatCh2.0 was more promising for all-optical applications: at 561 nm the photocurrent was only 2% and at 594 and 640 nm the photocurrent was undetectable (< 0.3%).

In some opsins, light at a red-shifted wavelength can reverse retinal isomerization, forcing the channel closed.^38,39^ To mimic the conditions of a typical all-optical electrophysiology experiment, we thus tested the combination of blue (240 mW cm^-2^) and intense orange (594 nm, 1 W cm^-2^) or red (640 nm, 8 W cm^-2^) light (Fig. 3I). The orange light had negligible activating or inactivating crosstalk into PsCatCh2.0 activation, but partially activated the ChR2-3M (^∼^20%). The red light slightly activated both constructs (^∼^10% for ChR2-3M and ^∼^5% for PsCatCh2.0), and also substantially inactivated ChR2-3M, leading to a ^∼^40% drop in photocurrent. Together, these results indicate that PsCatCh2.0 is a particularly promising channelrhodopsin for all-optical physiology experiments.

We then tested the new opsins in cultured rat hippocampal neurons (Fig. 4). Under paired optogenetic stimulation and pH imaging (Fig. 4A,B), we observed acidification in cells expressing CheRiff (pH = 6.87 ± 0.27, mean ± S.D., *n* = 24 cells), as before. We observed substantially less acidification in cells expressing either ChR2-3M (pH = 7.13 ± 0.19, mean ± S.D., *n* = 31 cells, *p* = 2.5e-4), or PsCatCh (pH = 7.14 ± 0.11, mean ± S.D., *n* = 25 cells, *p* = 4e-5, Fig. 4C-E). We then produced cultures co-expressing each of the three channelrhodopsins and QuasAr6a for voltage imaging (Fig. 4F,G). Under blue light stimulation, each opsin induced reliable spiking (Fig. 4H). ChR2-3M also induced some firing in intervals after a blue light pulse, presumably due to the very slow closing of the channel (τ_off_ = 1950 ± 500 ms, Fig. 3F) leading to residual currents.

**Figure 4.**
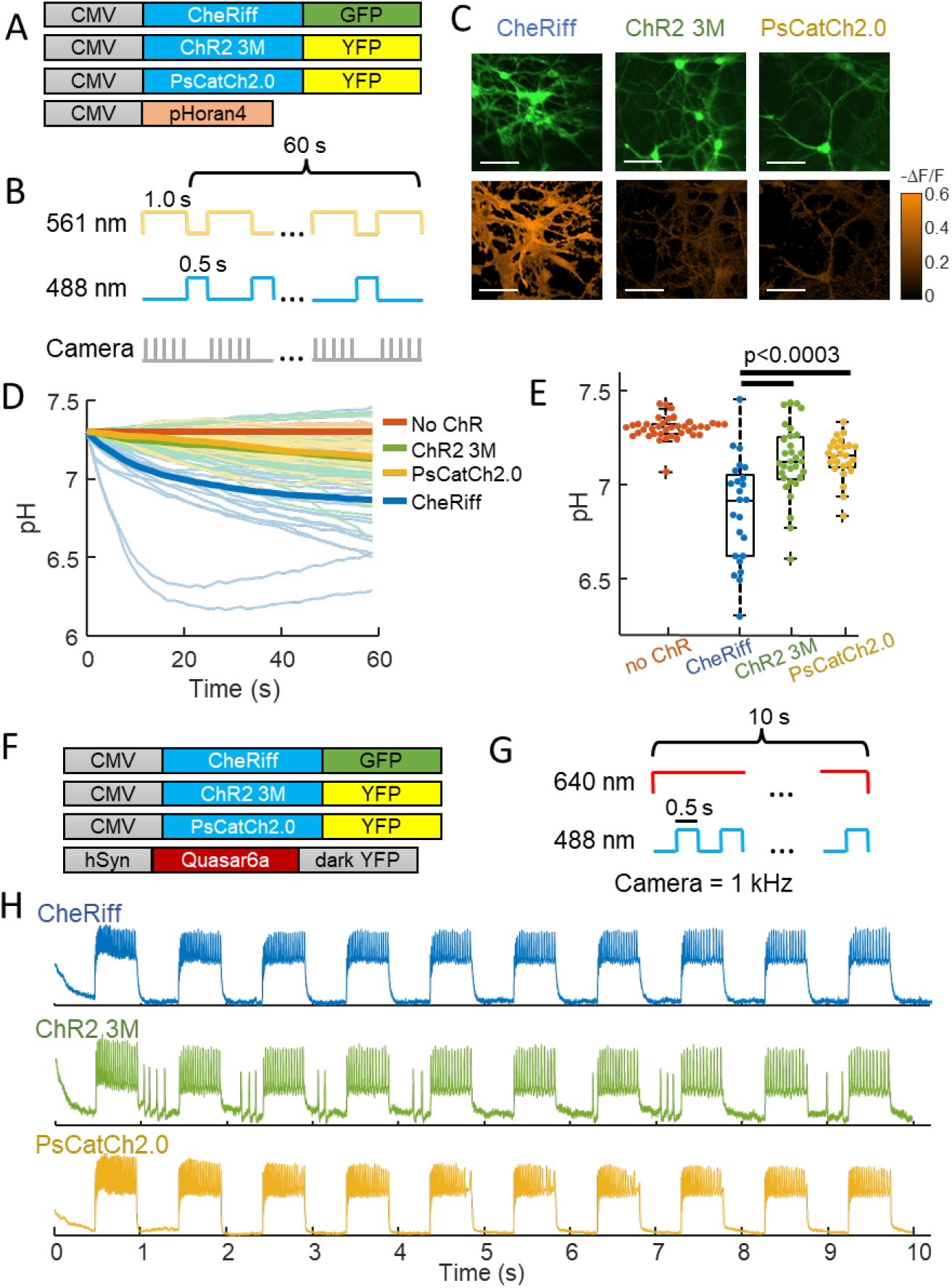
Chr2-3M and PsCatCh2.0 acidify neurons less than CheRiff. A) Genetic constructs for simultaneous optogenetic stimulation and pH imaging. B) Experimental paradigm for measuring pH responses to optogenetic stimulation. Stimulation (blue) and measurement (yellow) are interleaved for 60 s to avoid optical crosstalk. C) Images of cultured neurons showing (top) GFP or YFP fluorescence, a marker for Channelrhodopsin expression, (bottom) DF/F in the pHoran4 channel after the protocol shown in (B). (left) CheRiff-GFP, (middle) ChR2-3M-YFP, (right) PsCatCh2.0. Scale bars xx mm. D) Time-course of pH dynamics in cultured neurons. Cells expressing ChR2-3M and PsCatCh2.0 acidify less than CheRiff. E) Neurons expressing ChR2-3M, pH = 7.13 ± 0.19 (mean ± S.D., *n* = 31 cells), and PsCatCh2.0, pH = 7.14 ± 0.11 (mean ± S.D., *n* = 25 cells) had significantly less acidification (*p* = 2.5e-4, *p* = 4e-5, respectively, Wilcoxon signed-rank test) than CheRiff-expressing neurons, pH of 6.87 ± 0.27 (mean ± S.D., *n* = 24 cells). F) Genetic constructs for simultaneous optogenetic stimulation and Voltage imaging. G) Experimental paradigm for measuring voltage responses to optogenetic stimulation. Stimulation (blue) and measurement (red). H) Time-course of optogenetically activated spiking in cultured neuron expressing (top) CheRiff, (middle) ChR2-3M, or (bottom) PsCatCh2.0.

## Discussion

Most polarized cells have a strong inward-directed proton-motive force, and sodium-proton exchangers are required to maintain physiological intracellular pH.^40,41^ A sudden change in proton permeability of the membrane can disrupt this balance, leading to intracellular acidification. The proton flux was sufficient to overwhelm the buffering capacity of the cytoplasm. While we focused on the acidification due to CheRiff, we expect similar levels of acidification from other channelrhodopsins, unless they have been engineered specifically to be proton-impermeable.

The degree of acidification depends on the sub-cellular opsin distribution, the cell geometry, and the stimulus protocol. In general, the smaller the compartment, the larger the acidification. While we could not resolve individual dendritic spines, our results suggest that if channerhodopsins exist in spines, optogenetic acidification in spines could be substantial. In some all-optical physiology experiments, channelrhodopsins are restricted to the soma and proximal dendrites via trafficking sequences such as the K_v_2.1 trafficking motif.^42^ While this restriction has been primarily to facilitate targeted single-cell activation, a possible side effect is to decrease acidification in small distal compartments (dendrites, spines, axon).

Here we showed that the high-performance opsins ChR2-3M and PsCatCh2.0 have very low proton permeability, enabling repeated stimulation with minimal local acidification. We observed no activation-induced acidification in HEK cells (Figs. 3C,D), but we did observe a very slight acidification in neurons (Figs. 4D,E). We speculate that the activity-induced neuronal acidification was due to cell autonomous mechanism ^43–45^ as opposed to proton transport through the opsins.

While the ChR2-3M construct is not optimal for all-optical physiology experiments due to crosstalk with longer wavelength light, several features make it promising for prospective therapeutic applications. It has a very high photocurrent and high sensitivity, meaning that substantial modulation can be achieved with light at intensities < 10 mW/cm^2^. The slow closing of this construct could enable tonic activation with pulsed light, further decreasing the optical dose into the tissue. The positive reversal potential (+16.6 mV) further contributes to the ability of this channelrhodopsin to depolarize cells, even when the cells are already partially depolarized. Together with its low proton permeability, these attributes make ChR2-3M a good candidate for therapeutic applications requiring slowly varying changes in optogenetic drive. However, the slow kinetics of ChR2-3M may limit its use for basic science applications such as circuit mapping. PsCatCh2.0 is more suitable for applications requiring precisely timed spikes.

## Acknowledgments

We thank He Tian, Hillel Ori, Andrew Preecha and Shahinoor Begum for helpful discussions and technical assistance. The BeRST1 dye was provided by Evan Miller. This work was supported by a Vannevar Bush Faculty Fellowship, and NIH grants 1-R01-MH117042 and 1-R01-NS126043.

## Materials and Methods

### HEK293T cell culture

HEK293T cells were purchased from ATCC (CRL-3216) and validated by STR profiling. Mycoplasma testing was negative. Wild-type or engineered HEK293T cell lines were maintained at 37 °C, 5% CO_2_ in Dulbecco’s Modified Eagle Medium (DMEM) supplemented with 10% fetal bovine serum, 1% GlutaMax-I, penicillin (100 U/mL), streptomycin (100 µg/mL). For maintaining or expanding the cell culture, we used TC-treated culture dishes (Corning). For all imaging experiments, cells were plated on PDL-coated glass-bottomed dishes (Cellvis, Cat.# D35-14-1.5-N).

### Neuron culture

Primary E18 rat hippocampal neurons (fresh, never frozen, BrainBits #SDEHP) were dissociated following vendor protocols and plated in PDL-coated glass bottom dishes (Cellvis, Cat.# D35-14-1.5-N). Neurons (21k/cm^2^) were cocultured with primary rat glia (27k/cm^2^) to improve cell health and maturation.

### Lentivirus preparation

All the lentivirus preparations were made in house. HEK293T cells were co-transfected with the second-generation packaging plasmid psPAX2 (Addgene #12260), envelope plasmid VSV-G (Addgene #12259) and transfer plasmids at a ratio of 9:4:14. For small batches, 2.7 μg total plasmids for a small culture (300k cells in 35-mm dish) gave sufficient yield of lentivirus. Some viruses were concentrated using Lenti-X Concentrator (Takara Cat. # 631232) following vendor protocols and were concentrated 1/10. Quantities of virus used were quoted as non-concentrated amounts.

### Expression of optogenetic actuators and reporters

HEK293T cells were transduced at least 2 days before imaging with 50-200 µL of lentivirus encoding the desired Channelrhodopsin. Cell lines were created for stable expression of pHoran4, and of pHoran4 with Dox inducible K_ir_2.1-CFP, using fluorescence activated cell sorting (FACS) on cells that already had stable rtTA3 expression through antibiotic selection. K_ir_2.1 expression was induced 2 days before imaging by adding 1 µg/mL doxycycline, which was kept on the culture until time to image.

Neurons were transduced after 6-10 days in culture with 1) 200 µL lentivirus encoding pHoran4 driven by the CMV promoter, or 100-200 µL lentivirus encoding Quasar6a driven by the synapsin promoter and 2) 50-200 µL of the Channelrhodopsin variants, also driven by the CMV promoter. Functional imaging was performed after 14-20 days in culture.

### pH calibration

The pH response of pHoran4 was calibrated by changing the buffer pH stepwise from 6.4 to 7.3 (Figure 1 – figure supplement 1). To equilibrate the pH of the cytosol with the buffer pH, we added the K^+^/H^+^ exchanger nigericin at 14 μM. To prevent a [K^+^] gradient from driving a proton gradient, we used a high-potassium extracellular buffer (^22^,^23^). The buffer composition was (in mM): Good’s zwitterionic buffer 25, KCl 100, NaCl 38, CaCl_2_ 1.8, MgSO_4_ 0.8, NaH_2_ PO_4_ 0.9. The Good buffer, chosen based on its pK_a_ and effective buffering pH range, was MES for pH 6.4 and HEPES for pH 6.7–7.3. After perfusion of the buffers with different pH values, we waited 1 minute for the pH to equilibrate and recorded the steady state fluorescence for each cell. ΔF/F was calculated using the pH 7.3 as the baseline. The ΔF/F at each pH was then averaged across cells, and this average was fit with piecewise linear interpolation, which was used for converting ΔF/F to pH in subsequent data analysis.

### Sample preparation for imaging

Before optical stimulation and imaging, 35 mm dishes were washed with 1 mL PBS to remove residual culture medium, then filled with 2 mL extracellular (XC) buffer containing (in mM): 125 NaCl, 2.5 KCl, 2 CaCl_2_, 1 MgCl_2_, 15 HEPES, 25 glucose (pH 7.3). All imaging and electrophysiology were done using this XC buffer. For voltage imaging experiments in neurons, we added 10 μM NBQX, 20 μM Gabazine, 25 μM AP-V to block synaptic transmission.

BeRST1 was a gift from Evan Miller (Berkeley) and was used for voltage imaging in HEK cell monolayers. Cells were washed to remove culture medium and then incubated with 1-2 μM BeRST1 dye in XC buffer for 30 minutes. Immediately before imaging, samples were washed twice and immersed in XC buffer.

### Combined optogenetic stimulation and imaging

Experiments were conducted on a home-built inverted fluorescence microscope equipped with 405 nm, 488 nm, 532 nm, 561 nm, 594 nm, and 640 nm laser lines and a scientific complementary metal-oxide semiconductor (CMOS) camera (Hamamatsu ORCA-Flash 4.0). Beams from lasers were combined using dichroic mirrors and sent through an acousto-optic tunable filter (Gooch and Housego TF525-250-6-3-GH18A) for temporal modulation of intensity of each wavelength. The beams were then expanded and sent either to a DMD (Vialux, V-7000 UV, 9515) for spatial modulation or sent directly into the microscope (to avoid power losses associated with the DMD). The beams were focused onto the back-focal plane of a 60×/1.2-NA (numerical aperture) water-immersion objective (Olympus UIS2 UPlanSApo ×60/1.20W) or a 20×/0.75-NA objective (Olympus UIS2 UPlanSApo ×20/0.75). For Green and Yellow fluorescent protein, pHoran4, and QuasAr6a, fluorescence emission was separated from laser excitation using a dichroic mirror (488/561/633). Imaging of pHoran4 fluorescence was performed with 561 nm laser at illumination intensities of 100-200 mW cm^−2^. Imaging of QuasAr6a fluorescence was performed with 640 nm laser at an illumination intensity of 8 W cm^−2^. Stimulation of Channelrhodopsins was performed with 488 nm laser at an illumination intensity of 400– 800 mW cm^−2^.

### Electrophysiology

For patch clamp measurements, filamented glass micropipettes (WPI) were pulled to a resistance of 5–10 MΩ and filled with internal solution containing (in mM) 6 NaCl, 130 K-aspartate, 2 MgCl_2_, 5 CaCl_2_, 11 EGTA, and 10 HEPES (pH 7.2). The patch electrode was controlled with a low-noise patch clamp amplifier (either A-M Systems model 2400 or Axon Instruments MultiClamp 700B). Current traces were collected in voltage clamp mode. The collected electrophysiology data had a moving average filter applied to help reduce noise. The time constants were fit using single exponentials. In plots with multiple wavelengths of stimulation, the currents were normalized to the peak current for 488 nm stimulation.

### Wide-field imaging and patterning

Spatially resolved optical electrophysiology measurements were performed using a home-built upright ultra-wide-field microscope^24^ with a large field of view (4.6 × 4.6 mm^2^, with 2.25 × 2.25 μm^2^ pixel size in the sample plane) and high numerical aperture objective lens (Olympus MVPLAPO 2XC, NA 0.5). The fluorescence of BeRST1 was excited with a 639 nm laser (OptoEngine MLL-FN-639) at 100 mW cm^−2^, illuminating the sample from below at an oblique angle to minimize background autofluorescence. BeRST1 fluorescence was separated from scattered laser excitation via a dichroic beam splitter (Semrock Di01-R405/488/561/635-t3–60x85) and an emission filter (Semrock FF01-708/75–60-D). Images were collected at a 100 Hz frame rate on a Hamamatsu Orca Flash 4.2 scientific CMOS camera. Optogenetic stimulation was performed by exciting Channelrhodopsins with a blue LED (Thorlabs M470L3) with a maximum intensity of 400 mW cm^−2^.

### Measuring permeability of ChR2-3M

*Xenopus* oocytes were injected with cRNAs and maintained at 16 °C for 2 days in ND96 solution: 96 mM NaCl, 2 mM KCl, 1 mM CaCl_2_, 1 mM MgCl_2_, 10 mM HEPES, pH 7.4, and 50 μg/mL gentamycin. Two-electrode voltage-clamp was used for oocyte electrophysiology with TURBO TEC-05 amplifier from NPI (NPI electronics GmbH, Tamm, Germany). For current amplitude comparisons, photocurrents were measured in extracellular solution containing 110 mM NaCl, 5 mM KCl, 2 mM MgCl_2_, 2 mM BaCl_2_, 5 mM HEPES, pH 7.6; holding at -70 mV. Shifts in reversal potential, V_r_, were calculated by the reversal potential differences upon changing extracellular Na^+^ or K^+^ concentration from 120 mM (120 mM NaCl/KCl, 2 mM BaCl_2_, 5 mM HEPES, pH adjusted to 7.6 by N-Methyl-D-glucamine) to 1 mM (1 mM NaCl/KCl, 119 mM N-Methyl-D-glucamine, 2 mM BaCl_2_, 5 mM HEPES, pH adjusted to 7.6 by HCl). For all oocyte experiments, 473 nm laser at 5 mW/mm^2^ was used for illumination.

### Data Analysis

All data were processed and analyzed in MATLAB. For recordings with interleaved optogenetic stimulation and pH imaging, camera frames during stimulation were discarded, and frames during each period of imaging were averaged. Baseline fluorescence, F_0_, was calculated from the first frame, before any optogenetic stimulation. A threshold on F_0_ was set to restrict calculation of ΔF/F_0_ to signal-bearing regions of the sample.

Individual cells expressing the desired constructs were selected and fluorescence waveforms were calculated by averaging pixels whose baseline value exceeded the threshold. Sensor photoactivation artifacts were characterized using matched controls that expressed pHoran4 but no channelrhodopsin. Population-average photoartifacts were subtracted from the signals obtained from cells with channerhodopsin expression.

Dendrites were selected and analyzed in the same manner as somas, and were then associated with the connected soma. Acidification half-times were calculated by finding the maximum acidification during the stimulation period, and then finding the time point where the ΔF/F first reached half of the maximum decrease. Recovery after stimulation was fit to a single exponential, and the fit function was used to calculate the half-recovery time. Statistical tests were done using the Wilcoxon signed-rank test.

## Materials Availability Statement

Plasmids developed for this study are available on Addgene: https://www.addgene.org/browse/article/28228806/

## Data Availability Statement

Data underlying each figure panel are available on Figshare at: https://figshare.com/projects/Diminishing_neuronal_acidification_by_channelrhodopsins_with_low_proton_conduction/178173

## Figures

**Figure 1 - figure supplement 1.**
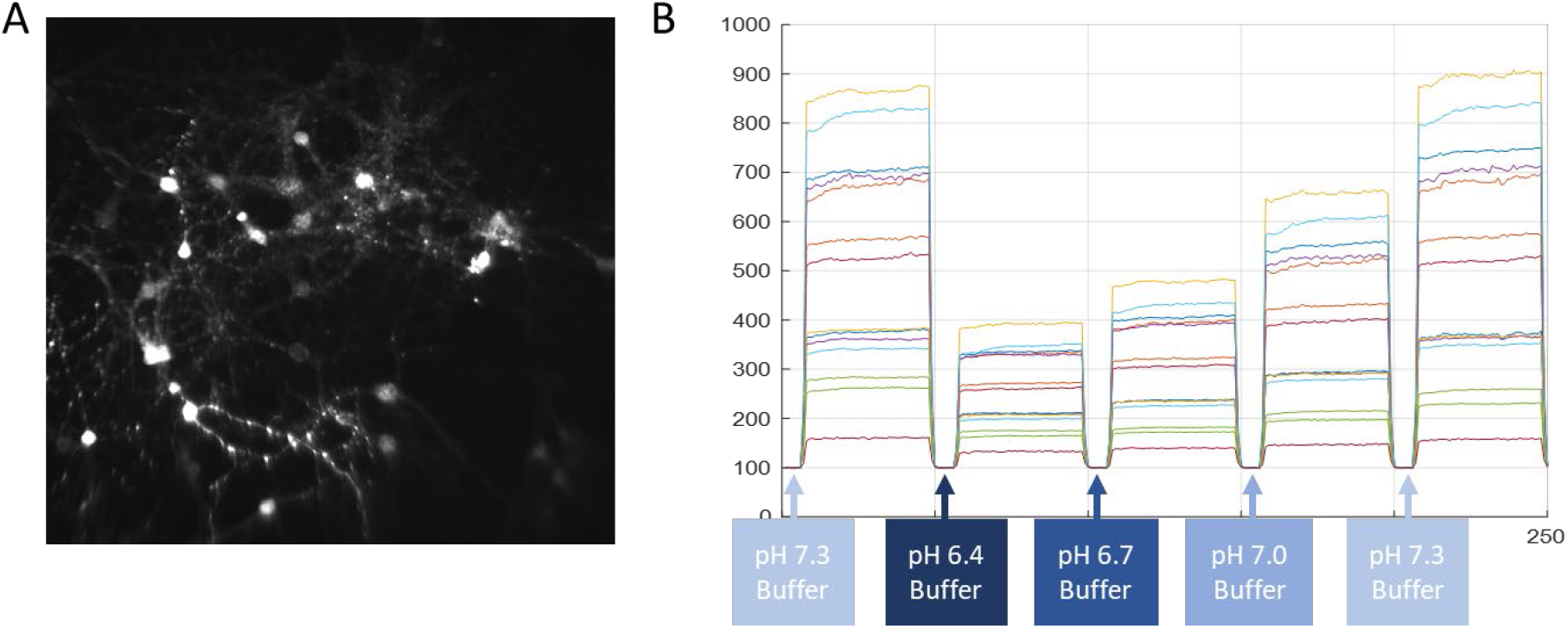
Procedure for calibrating pHoran4 pH measurements. A) Representative image of cultured neurons expressing pHoran4. The cells have been permeabilized with Nigericin and are in a high K^+^ extracellular medium (Methods). B) Example fluorescence traces of individual cells as the dish is perfused with buffers of different pH values.

**Figure 3 – figure supplement 1.**
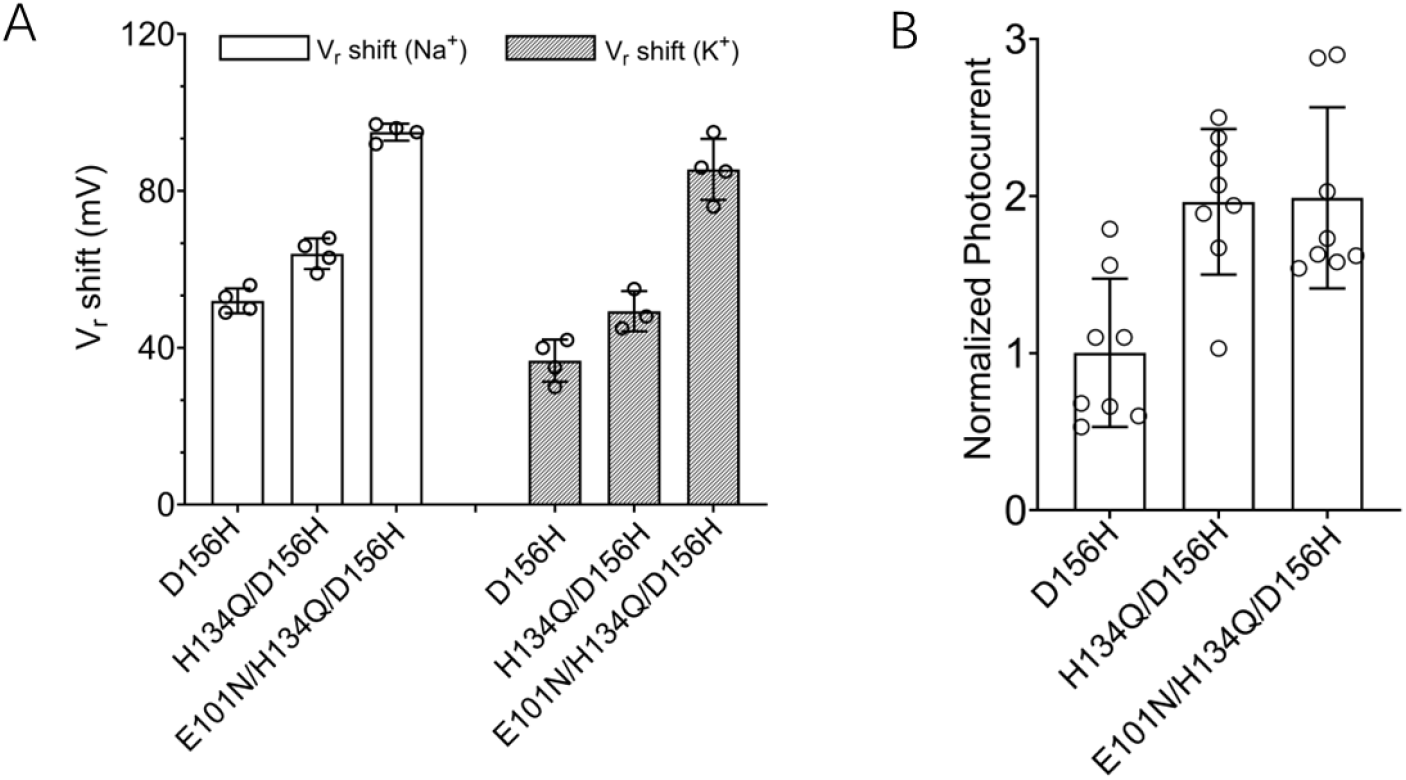
Engineering of ChR2-3M, a channelrhodopsin with high Na^+^ and K^+^ selectivity and high photocurrent amplitude. (A) Shifts in reversal potential (V_r_) of ChR2 variants upon changing extracellular Na^+^ or K^+^ concentration from 120 mM to 1 mM (mean ± S.D., *n* = 3-4 cells). (B) Photocurrent amplitudes of ChR2 variants (mean ± S.D., *n* = 8 cells). The triple mutant E101N/H134Q/D156H was modified with trafficking, ER export, and signal peptides and designated ChR2-3M. Experiments were performed with *Xenopus* oocytes expressing different Channelrhodopsin variants.

**Figure 3 – figure supplement 2.**
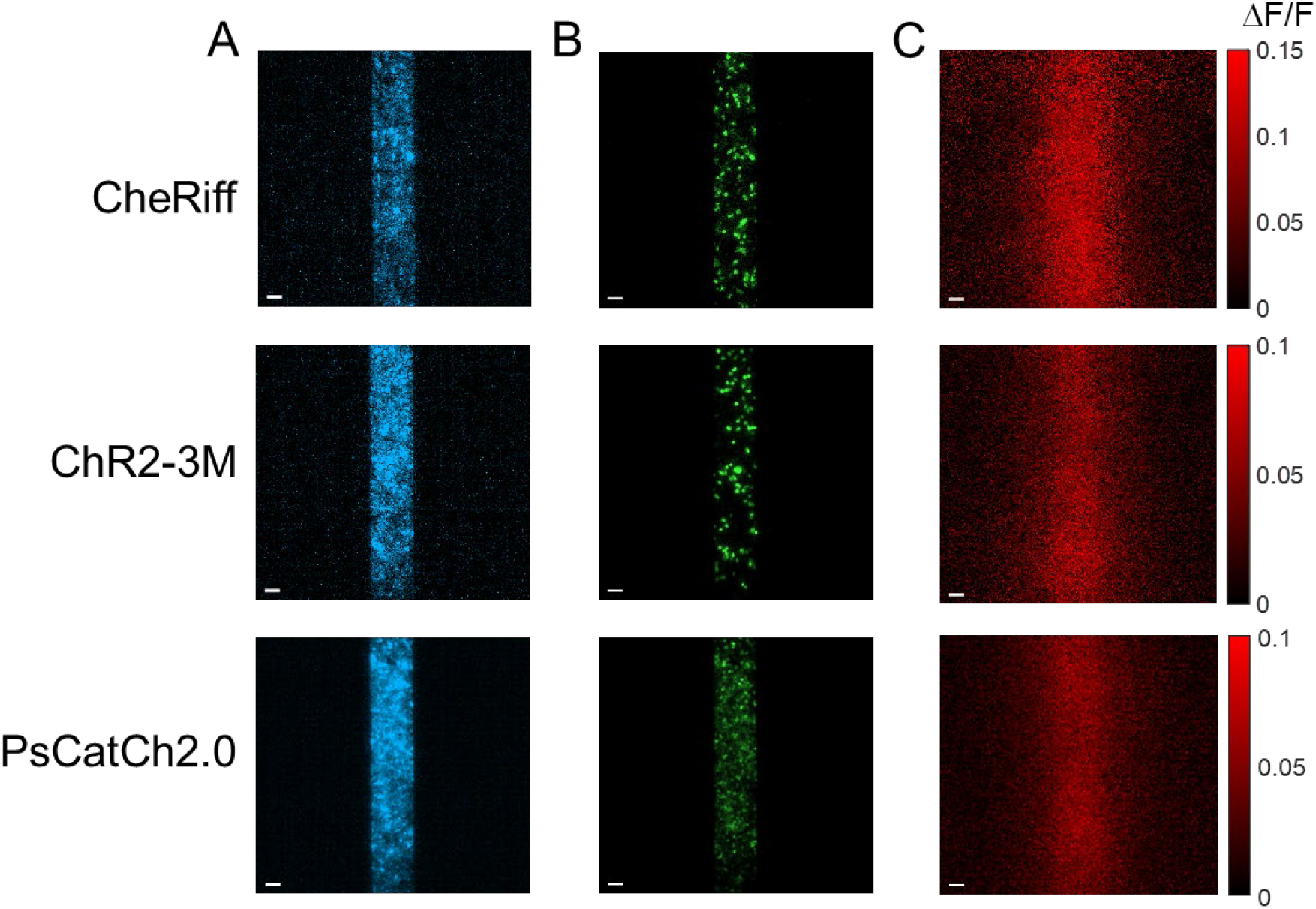
Depolarization of HEK cell monolayers via patterned stimulation of channelrhodopsins. (A) Blue light stimulation patterns. (B) Fluorescence of GFP or YFP tags on the opsins expressed in confluent monolayers of HEK cells, under patterned blue light excitation. (C) BeRST1 ΔF/F showing depolarization from patterned stimulation. Scale bars 100 µm.

